# Fixation probability in a diploid sexually reproducing population

**DOI:** 10.1101/2024.10.03.616511

**Authors:** Zhenyu Shi, Loïc Marrec, Xiang-Yi Li Richter

**Affiliations:** Department of Automation, Shanghai Jiao Tong University, Dongchuan Road 800, Shanghai, 200240, China; Institute of Ecology and Evolution, University of Bern, Baltzerstrasse 6, Bern, 3012, Switzerland; Department of Biology, University of Konstanz, Universitätsstraße 10, Konstanz, 78464, Germany

**Author notes:** Contributing authors.

**Keywords:** Birth-death process, Demographic stochasticity, Fixation probability, Sex ratio, Sexual reproduction, Sex-specific dominance coefficient

## Abstract

Classical population genetics models with haploid individuals and cloning reproduction, such as the Moran process, have limited power in predicting the evolutionary fate of mutants in diploid, sexually reproducing populations. In this work, we built a new stochastic population genetics model to fill this outstanding knowledge gap. We showed that sexual reproduction tends to suppress the fixation probability of beneficial mutants than clonal reproduction, except under overdominance. This effect is particularly large in small populations with biased sex ratios. We also found that small populations are prone to the invasion of mildly detrimental alleles when the heterozygotes have a fitness advantage. Our study highlights the pressing need to extend the research of stochastic population dynamics to consider the biology of sexual reproduction, such as allelic interactions and biased sex ratio. Our model extends the frontier of population genetics research into diploid, sexually reproducing populations and sets the foundation to explore the great variety of biological factors and processes, such as sexual selection and sexual conflict, on stochastic evolutionary dynamics. Future research in this direction will help make better predictions of the evolutionary fate of mutants in relevant contexts, including the conservation of endangered species and evolutionary rescue.

## 1 Introduction

The fixation probability is the probability that an allele reaches a frequency of one in a population, which makes it a fundamental concept of evolution [1]. In particular, the fixation probability allows for assessing whether a population is likely to adapt to changing environments and predicting the rate of loss of genetic diversity. For example, one can use the fixation probability to assess the risk of resistant bacteria taking over a bacterial population during antibiotic treatment, which is important for public health [2]. In conservation biology, the fixation probability may help determine whether beneficial mutations can counter the human-induced demographic decline of animal populations, thus contributing to population management strategies [3].

The variety of applications of the fixation probability explains the interest in calculating it, which dates back to the work of Fisher and Haldane [4–7]. Many methods exist to calculate the fixation probability, such as based on branching processes or diffusion approximations [1]. Another method is based on modeling the evolutionary dynamics of a population as a Markov chain and computing the probability of reaching an absorbing state [8]. This is the case with the Moran process, in which at each time step, one haploid individual reproduces and another dies until the fixed-size population becomes monomorphic [9].

In the simple case of a well-mixed population, the Moran process allows one to derive an equation for the fixation probability [10]. Since most populations in the wild harbors some spatial structure, this property was later included in the Moran process, giving rise to Evolutionary Graph Theory [11]. This theory aims to quantify how spatial structure impacts the fixation probability. It showed that some spatial structures amplify natural selection, i.e., increase the fixation probability of beneficial mutations and decrease that of deleterious mutations compared to well-mixed populations, whereas others suppress natural selection [12].

Up to now, sexual reproduction has not been considered in most population genetics models. For example, the Moran process has so far been confined to haploid individuals reproducing asexually. However, diploidy and sexual reproduction have the potential to profoundly impact the evolutionary dynamics of populations. For example, the phenotype and fitness of a heterozygote can be intermediate between the mutant and ancestral homozygotes, or exceed this range under overdominance (positive heterosis) or underdominance (negative heterosis) [13]. The dominance coefficient of alleles can also differ between the sexes [14]. Such flexibility allows a variety of asymmetry to arise between the sexes. For example, phenotypes corresponding to the same genotype can differ between sexes (sexual dimorphism) and have different fitness effects [15]. Such intralocus sexual conflict can be ameliorated by sex-specific dominance reversal, where the female/male heterozygote resembles the phenotype of the homozygote of the female-/male-beneficial allele, respectively [16, 17]. In addition, the sex ratio of populations can deviate from being balanced due to intrinsic (e.g. maternal manipulation) and external (e.g. selective hunting) causes, and the diversity of mating systems (e.g., monogamy and polygyny) can cause both the intensity and softness of selection to differ between the sexes [18]. These asymmetries make it difficult to predict the evolutionary fate of mutants.

Therefore, in the current work, we extend the classical Moran process to diploid individuals and sexual reproduction. Specifically, we aim to compute the fixation probability of a mutant allele and, thus, determine whether sexual reproduction amplifies or suppresses natural selection and under which conditions. We consider that amplification (respectively suppression) of natural selection occurs if the fixation probability of a beneficial allele is greater (respectively lower) than that under the classical Moran process in a population of the same size (i.e., number of alleles). We start from computing the fixation probability of a mutant allele in the basic scenario with a balanced sex ratio and identical dominance coefficients in males and females. Then, we quantify how varying the sex ratio and sex-specific dominance coefficients can impact the fixation probability.

## 2 Results

### 2.1 An extension of the Moran process with diploid individuals and sexual reproduction

We consider a fixed-size population of *N*_F_ females and *N*_M_ males. Each individual has a diploid genetic locus with two different variants of alleles (i.e., A: mutant; B: ancestral). There are thus three possible genotypes: AA, AB, and BB. We assume their relative fitness to be 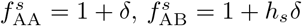, and 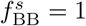, where *δ* is the fitness effect of the mutant, *h* is the dominance coefficient, and *s* ∈ {M, F} stands for the biological sex of the individual. Here, the fitness of the heterozygote can differ between the sexes. Similar to the Moran process, the population dynamics can be captured by a discrete-time birth-death process (Figure 1). In each time step, a male and a female are chosen to reproduce, with probabilities proportional to their fitness. Their gametes randomly combine to produce a diploid offspring. The sex of the offspring is chosen randomly with equal probability. The offspring will then replace a randomly chosen adult of the same sex. The process continues until the population becomes monomorphic (i.e., one of the alleles disappears from the gene pool).

**Fig. 1.**
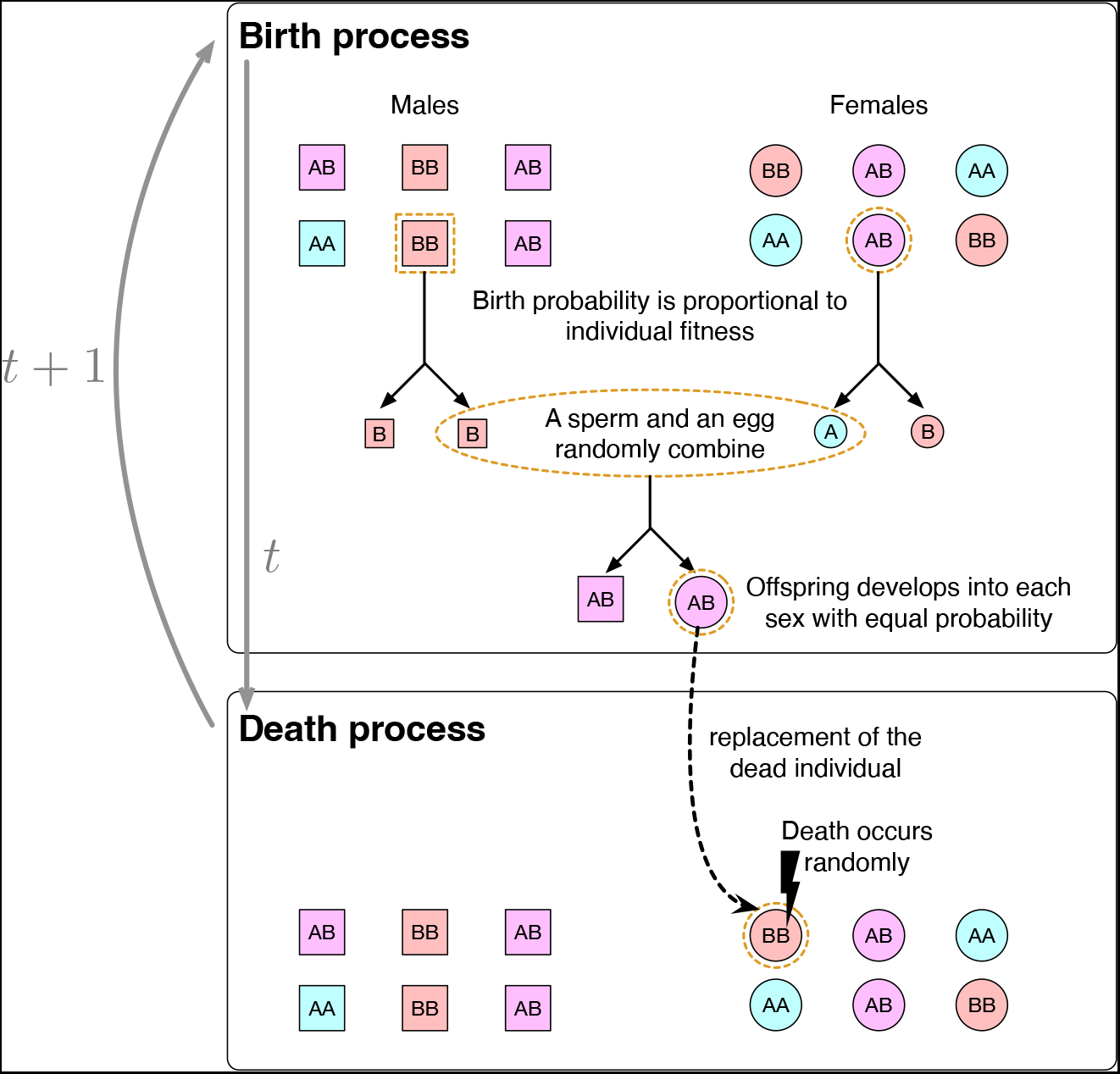
An illustration of the birth-death process with diploid sexually reproducing individuals.

The birth-death process can be conveniently adapted to model the sexual reproduction between monoecious individuals by replacing males and females with hermaphrodites.

### 2.2 Sexual reproduction generally suppresses natural selection, except under overdominance

To study the evolutionary fate of a newly arisen mutant allele (i.e., occurring in an AB heterozygote) in an ancestral population of BB individuals, we start from the basic case in a population with equal numbers of males and females (*N*_M_ = *N*_F_ = *N*). We also assume that the dominance coefficients of the mutant allele to be identical for both sexes (*h*_M_ = *h*_F_ = *h*). The fixation probability, *ρ*, can be calculated numerically and by simulations (see Methods). When the fitness effect of the mutant allele is small (*δ* → 0), we can use weak selection methods to calculate the selection gradient *ρ*^*′*^(0) analytically (see SI Appendix, section 2). Denoting the selection gradient as *µ*, in a *dioecious* population, we obtain

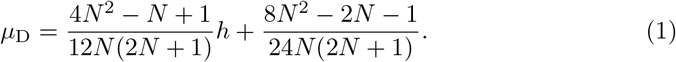

We also calculated the selection gradient of a single mutant allele in a population of *N* diploid hermaphrodites under weak selection, which has the same expression as *µ*_D_ in Eq.(1). However, to make the cases comparable, we must keep the total number of alleles identical. Therefore, we calculated the corresponding selection gradient in a population of 2*N monoecious* individuals, which is given by

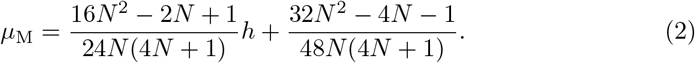

In large populations (*N* → ∞), the selection gradient of the mutant allele increases approximately linearly with the dominance coefficient of the mutant allele:

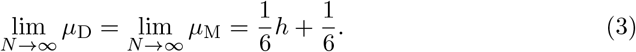

Although *µ*_M_ and *µ*_D_ have the same limit when the population size approaches infinity, *µ*_M_ is always greater than *µ*_D_. This is demonstrated in Figure 2a-b – given the same *h* value, the slope of the solid line at *δ* = 0 is always greater than that of the broken line when *h* is positive, and the difference is more pronounced in smaller populations.

**Fig. 2.**
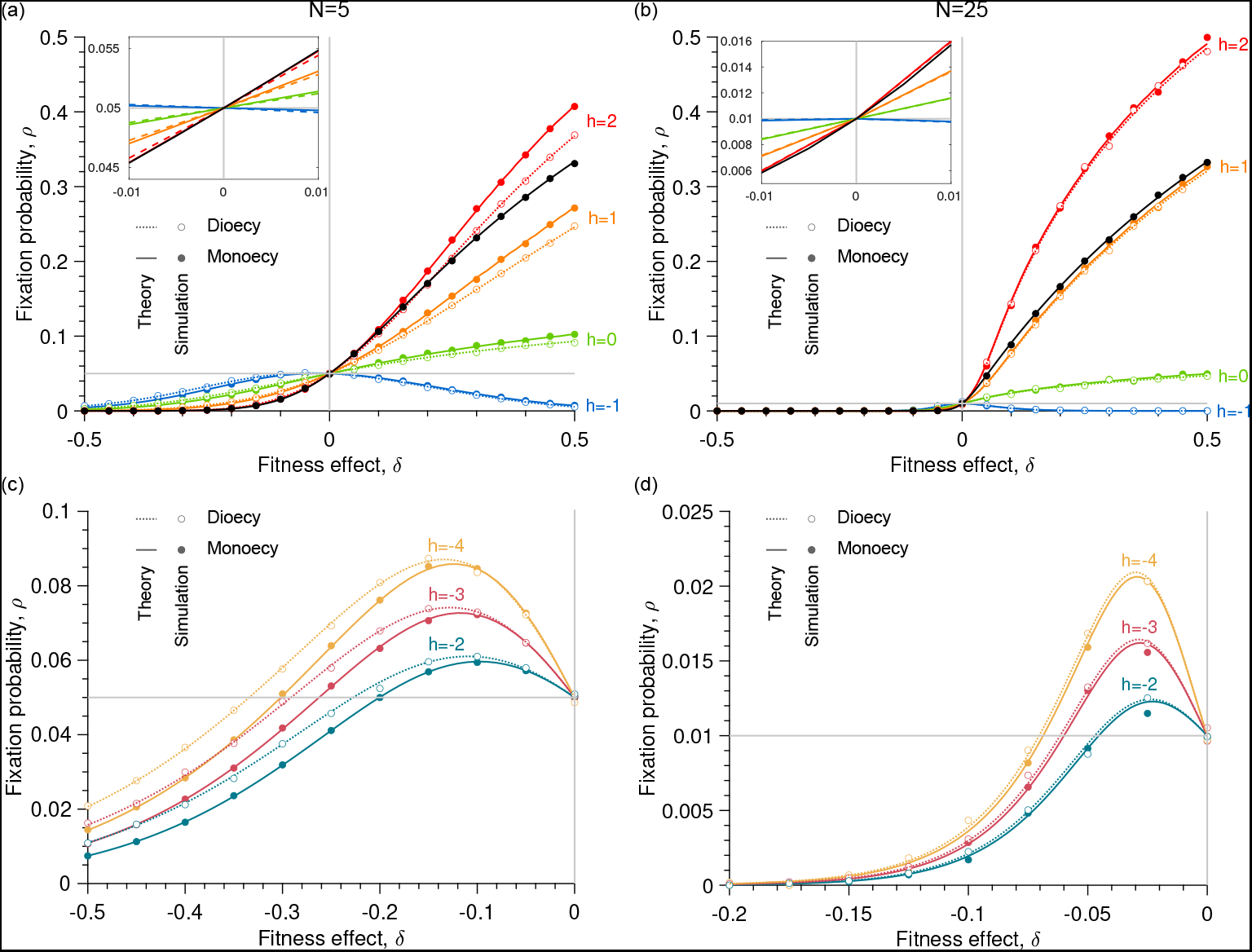
The fixation probability of a single mutant allele can be influenced by the population size (*N*), its fitness effect (*δ*), its dominance coefficient (*h*), and the mode of sexual reproduction (i.e., monoecy vs dioecy). The thick black line represents the fixation probability of the mutant allele under the classical Moran process in a population of size 4*N* (Eq. 4). The insets show the results under weak selection (*δ* values in the vicinity of zero). Each data point was averaged over 10^4^-10^5^ stochastic replicates.

In the classic Moran process, individuals are haploid. Therefore, to keep the total number of alleles identical to a diploid monoecious or dioecious population of 2*N* individuals, we need to use the fixation probability of a single mutant allele in a population of 4*N* haploid individuals,

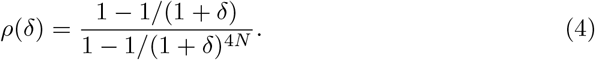

By Taylor expansion, we have the selection gradient of a single mutant allele at *δ* = 0,

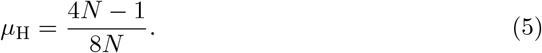

In large populations, we have

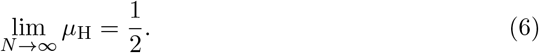

A comparison between (Eq. 3) and (Eq. 6) shows that if the fitness of the heterozygote is between the ancestral and mutant homozygotes (*h* ∈ [0, 1]), the selection gradient of the mutant allele is always smaller under sexual reproduction, no matter whether individuals are monoecious or dioecious (Figure 2a-b).

However, sexual selection does not always suppress natural selection. Under over-dominance (*h >* 1, the fitness of the heterozygote being greater than that of the homozygotes for beneficial mutants), the selection gradient at *δ* = 0 under sexual reproduction can exceed that of the classical Moran process. Our analysis shows that this occurs when the dominance coefficient is greater than 2 (SI Appendix, section 3.1).

The case of underdominance (*h <* 0, the fitness of the heterozygote being lower than both homozygotes for beneficial mutants), also produces interesting results. For example, when *h* = −1, the selection gradient at *δ* = 0 approaches zero when *N* → ∞ (SI Appendix, section 3.1). This means that in large populations, the fixation probability of beneficial mutants can rarely exceed the neutral expectation, and the fitter the mutant homozygote is, the less likely that it can reach fixation (Figure 2a-b, blue lines). However, when *h* is strongly negative, the fixation probability of mildly deleterious mutants can exceed the neutral expectation, and the range of fitness effect where this can happen is broader in smaller populations (Figure 2c-d). This means that smaller populations are more vulnerable to the invasion of deleterious mutations under sexual reproduction, if the heterozygote has higher fitness than the ancestral homozygote.

### 2.3 Biased sex ratio suppresses natural selection in small populations

In sexually reproducing populations with dioecious individuals, the sex ratio may not always be balanced. Up to now, the sole difference between the sexes is their label, therefore, without loss of generality, we only analyze the case where the mutant allele occurs in a female heterozygote. Here, we consider a population with a total of 2*N* individuals, while *N*_M_ = *kN*_F_. We also keep the assumption of *h*_M_ = *h*_F_ = *h* for now. In this case, the selection gradient at *δ* = 0 can also be derived analytically (SI Appendix, section 2.4). In large populations, biased sex ratios have negligible effects on the selection gradient (i.e., the sex ratio *k* is absent in Eq. (7)):

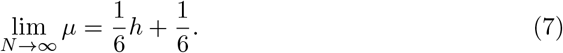

However, in small populations, biased sex ratio generally suppresses natural selection when the dominance coefficient is positive, and the effect increases with the value of *h* (Figure 3a-b). However, when *h* is sufficiently small, a beneficial mutant (i.e., with fitness effect *δ* greater than 0) can reach fixation more easily in populations with biased sex ratios (Figure 3c-d).

**Fig. 3.**
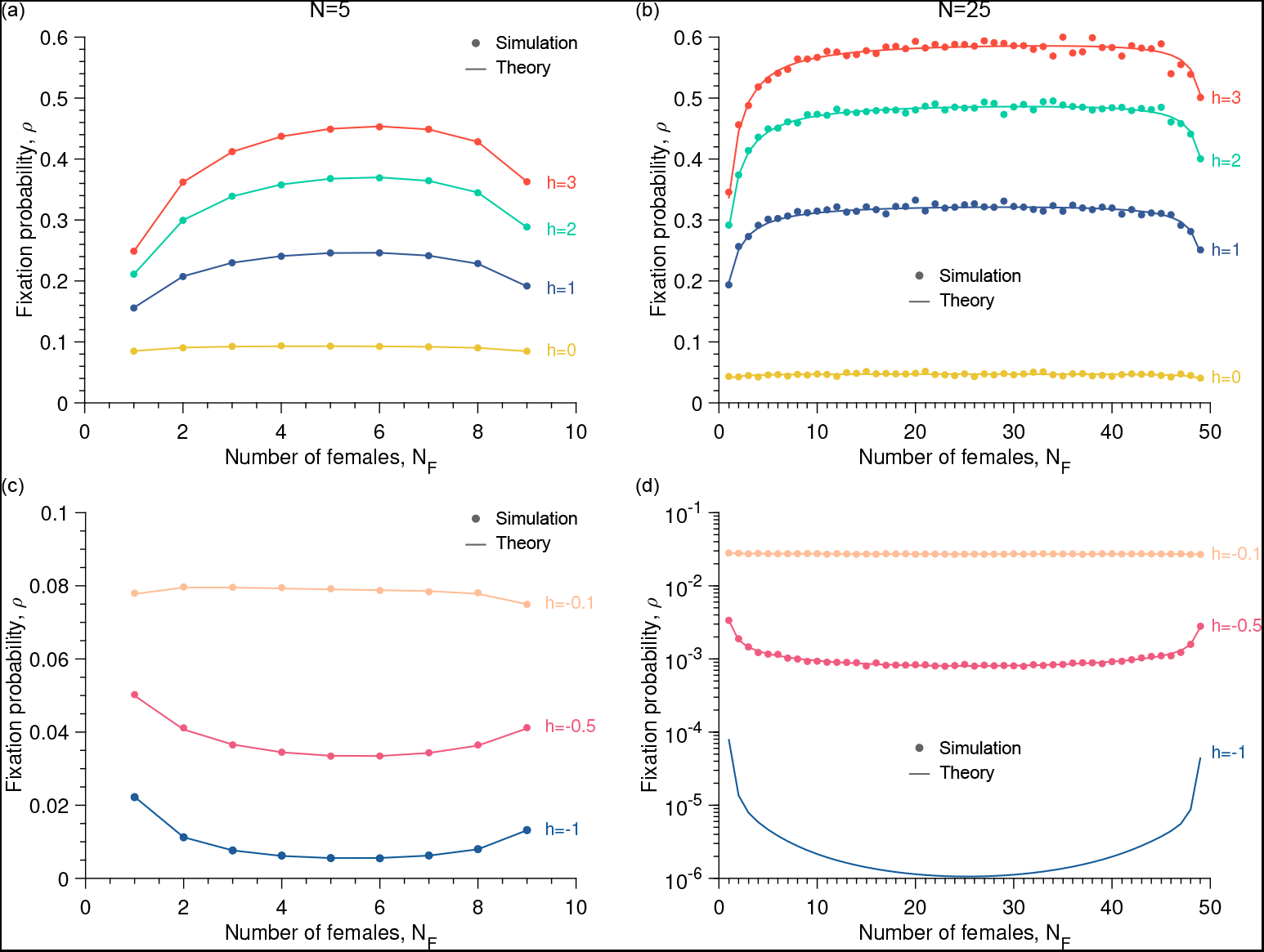
The dominance coefficient (*h*) of the mutant allele and the population’s sex ratio interact to influence the fixation probability of the mutant allele. The fitness effect (*δ*) of the mutant allele was set to 0.5. Each data point was averages over 10^6^ stochastic replicates.

### 2.4 Sex ratio and sex-specific dominance coefficients interact to influence the fixation probability of mutants

In dioecious species, the dominance coefficient of mutants may also differ between sexes. In this case, the fixation probability of the mutant allele can be influenced not only by the sex ratio of the population but also by the sex of the heterozygote where it first arises. The selection gradient *µ* at *δ* = 0 can still be calculated analytically under weak selection (SI Appendix, section 2.4). When the population size is sufficiently large, it is determined solely by the sex-specific dominance coefficients, *h*_M_ and *h*_F_, and not by the sex ratio:

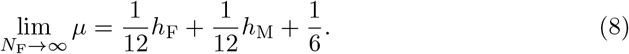

In small populations, however, sex ratio of the population can strongly influence the fixation probability of the mutant. Assuming that the mutant allele first arises in a female heterozygote, if it is dominant in females but recessive in males (*h*_F_ = 1 and *h*_M_ = 0), the fixation probability of this allele increases as the sex ratio of the population becomes more female-biased. However, if the mutant is dominant in males but recessive in females (*h*_M_ = 1 and *h*_F_ = 0), its fixation probability decreases as the sex ratio of the population becomes more female-biased (Figure 4a-b).

**Fig. 4.**
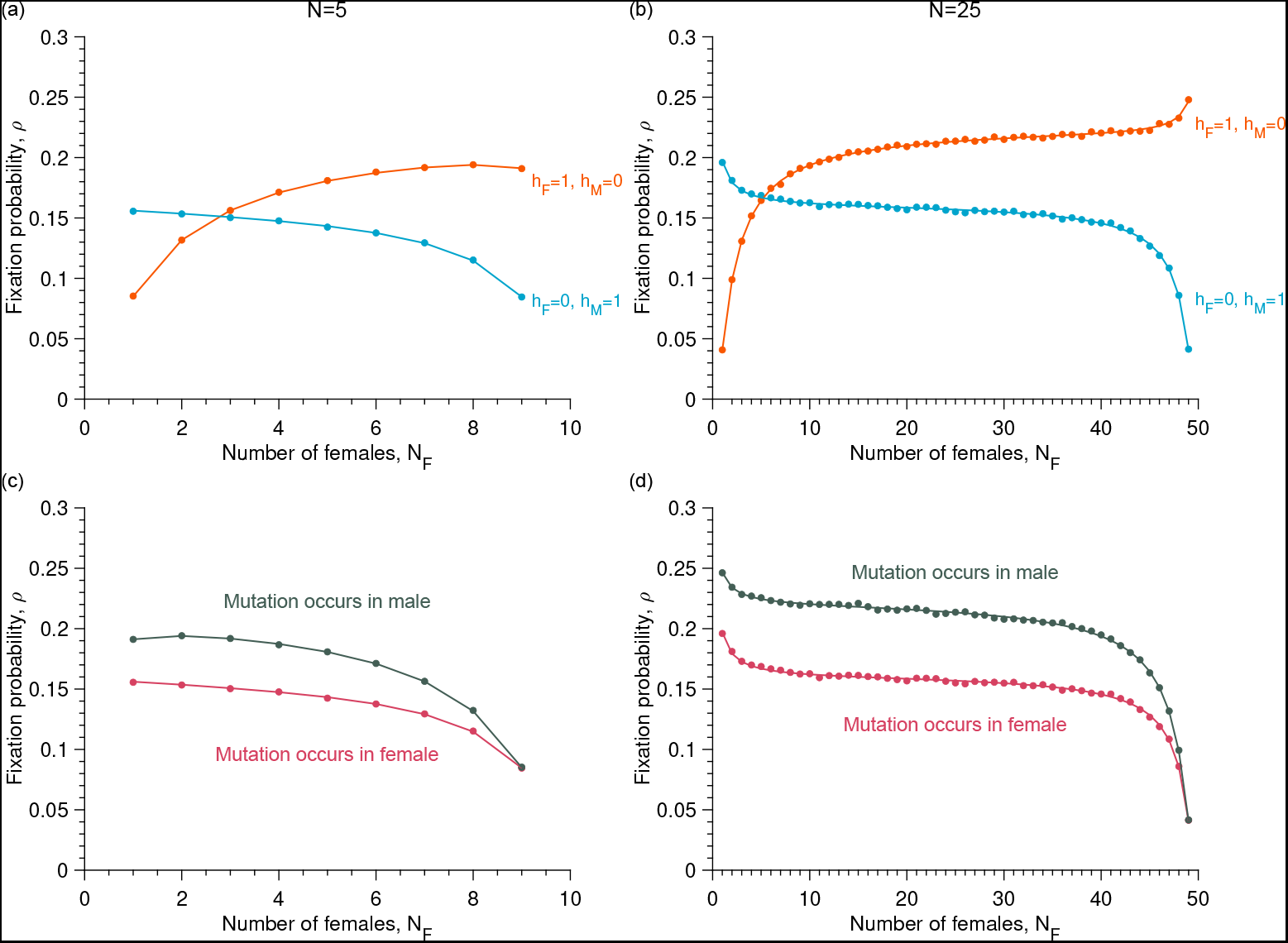
The fixation probability of a beneficial mutant allele (*δ* = 0.5) is influenced by the population sex ratio and the sex-specific dominance coefficients. In panels (c) and (d), the mutant allele is dominant in males and recessive in females. Each data point was averaged over 10^5^-10^6^ stochastic replicates.

Given the same total population size 2*N*, regardless of the sex ratio, a beneficial mutant always has higher fixation probability if it arises in the sex with higher dominance coefficient. For example, if the mutant is dominant in males but recessive in females, its fixation probability is always higher if the first heterozygote is male, independent of the sex ratio (Figure 4c-d).

## 3 Discussion

In this work, we developed a stochastic population genetics model with diploid, sexually reproducing individuals. In each time step, an individual is born to replace an adult to keep population size constant, resembling the classical Moran process. Despite being highly idealized, our model provides an opportunity to study stochastic evolutionary dynamics of mutants under sexual reproduction between monoecious or dioecious individuals. In the latter case, we also evaluated the effect of biased population sex ratio and sex-specific dominance coefficients of the mutants.

We showed that sexual reproduction generally suppresses natural selection, except under overdominance (Figure 2a-b). This effect is particularly strong under biased sex ratios in small populations (Figure 3). In other words, sexual reproduction and biased sex ratios increase the fixation probability of deleterious mutations and decrease that of beneficial mutations compared to a well-mixed haploid population of the same number of alleles. Thus, the fixation-effective population size *N*_e_ of such diploid populations is likely to be lower than their census size [19], which is in agreement with previous results [20] The fixation probability, as well as the effective size, is often derived from a diffusion approximation that consists of calculating the expectation and variance of changes in allele frequencies. We applied this method to reproduce the fixation probability of mutants in large monoecious populations, which confirmed our numerical and simulation results (SI Appendix, section 4). Furthermore, we were able to explore conditions that are not applicable for the diffusion approximation method, such as small population size and complex allelic interactions. Our study showed interesting and surprising results in such situations, for example, deleterious mutants (*δ <* 0) can have a fixation probability higher than the neutral expectation under negative heterosis (*h <* 0) (Figure 2c-d). In this case, the fitness advantage of the heterozygote contributes to the fixation of the mutant allele, although this will eventually cause a decrease of the fitness of the entire population once the deleterious allele reaches fixation. These cases are particularly relevant in the conservation of endangered species and ecosystem restoration, given that most charismatic keystone species and umbrella species are diploid and reproduce sexually.

We developed the model as a proof-of-principle to illustrate the pressing need of incorporating sexual reproduction into stochastic population dynamics models. In its current form, the model does not capture the influence of several important biological factors and processes such as frequency-dependent selection and fluctuating population size. We can overcome these limitations by extending the model using individual-based simulations. Our simulations showed excellent agreement with the theoretical results, and were fast to run. The simulation code (Supplementary Information) can serve as the basis for more complex models in the future.

Despite the unlimited possibilities of extending the current model for biological realism, we suggest prioritizing future research to explore the following directions, which we find the most interesting. The first is biased sex ratio caused by demographic stochasticity and population extinction due to the disappearance of one of the sexes. In small populations, it is interesting to explore the time scale for this to occur in comparison with the time scale of the fixation of mutants (e.g., in the context of evolutionary rescue). Another interesting topic to explore is the impact of sexually antagonistic alleles on stochastic evolutionary dynamics. Existing theoretical models investigating the impact of sexual antagonism on the maintenance of genetic variance generally ignore demographic stochasticity [17, 21–24], while empirical research on the evolutionary consequences of sexual antagonism, especially those involving experimental evolution, are often performed in populations of small sizes [25–28]. Solving this mismatch requires developing stochastic population dynamics models. Furthermore, the fitness of genotypes are often not fixed, as assumed in the current model, but can vary according to the natural and social environments [29–31]. Therefore, it would be very interesting to incorporate frequency-dependent selection, for example, by introducing spatial heterogeneity and dispersal, or fitness determination through evolutionary games.

In conclusion, our work shows the profound impact of the biology of sexual reproduction on stochastic population dynamics. Being vastly diverse and highly conserved across the eukaryotic domain, sexual reproduction has been a ubiquitous and indispensable component of evolution that produced the biodiversity of today’s world. Facing the current anthropogenic biodiversity crisis, there is an urgent need to investigate the evolutionary dynamics of sexually reproducing populations to help develop conservation and population management strategies. The model and methods we developed in this paper enable us to evaluate the influences of relevant biological factors, such as sex ratio and sex-specific dominance coefficients of alleles, on the evolutionary trajectory of small populations. We are now in a position to make better predictions of the population dynamics of sexually reproducing species by considering the role of demographic stochasticity, bringing us closer to the ultimate goal of reversing the trend of biodiversity loss and restoring the earth’s ecosystems.

## 4 Methods

### 4.1 Numerical solution of the fixation probability

To calculate the fixation probability of a mutant, we first define the states of the population and calculate the transition probabilities between all pairs of states in a birth-death cycle. In a population of *N*_F_ females and *N*_M_ males, the number of individuals of genotype *i* and sex *s* is denoted as 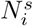 We denote the state of the population by a 4-tuple 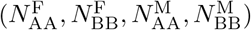 In each step, the probability that an individual of genotype *i* and sex *s* is selected to give birth is

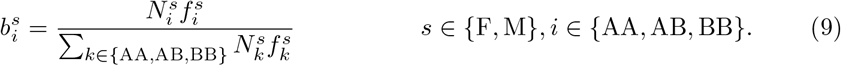

Assuming fair segregation of alleles during meiosis (i.e., alleles A and B enter the gametes of a heterozygote with equal probability) and random fusion between male and female gametes, the probabilities of the different alleles entering a new born individual are given by

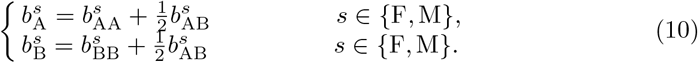

The probabilities that the new born offspring is of genotype AA, AB, or BB are thus

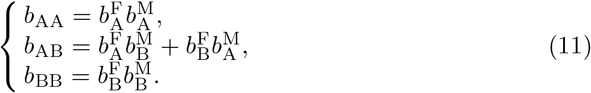

Assuming that death occur randomly, the probabilities of an individual of each genotype being replaced are given by

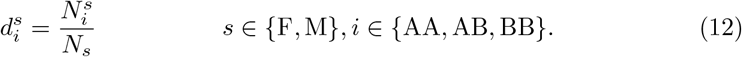

Based on these, the transition probabilities between two different population states in a birth-death cycle are given by

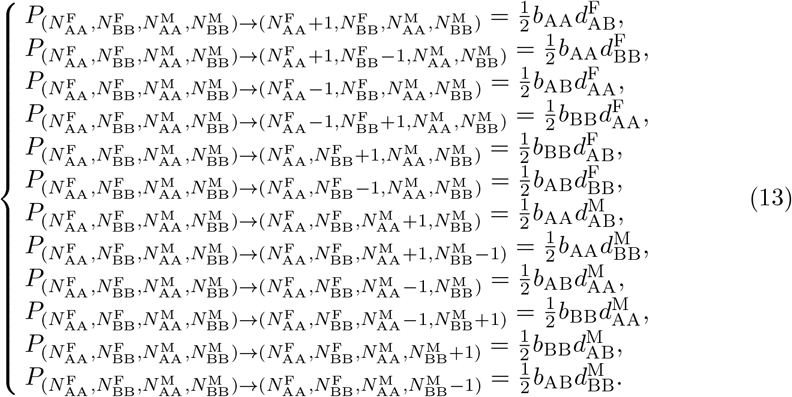

The sum of the above 12 probabilities is denoted as *P* 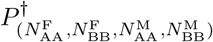 and the probability that a state stays unchanged is

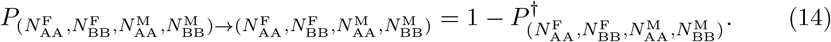

The transition probabilities between all other pairs of states are 0.

Starting from any initial state, the population will eventually become homogeneous. We denote the absorbing state of all individuals being AA or BB as **A** and **B**, respectively. Given a specific initial state **x**, the probability that it reaches the absorbing state **A** is denoted as *ρ*_x_, which is the so-called fixation probability. The fixation probabilities of different population states satisfy

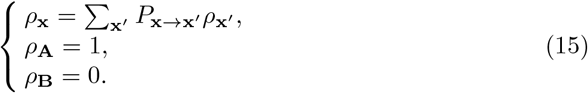

The above linear system of equations has a unique solution, and the time complexity of the solution is 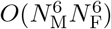

We focus on studying the fixation probability of a single mutant allele A occuring in an AB heterozygote in an ancestral population of BB individuals. Because of the symmetry between the two sexes (except when *h*_M_ ≠ *h*_F_), the fixation probability of the mutant allele does not depend on the sex of the first heterozygote. Therefore, without otherwise specified, we assume that the first heterozygote to be female and denote the fixation probability of from this state *ρ* (0,*N*_F_*−*1,0,*N*_M_) as *ρ* for convenience.

### 4.2 Analysis of the selection gradient by weak selection approximation

In general, the fixation probability can only be solved numerically. However, when the fitness effect *δ* is very small, we can perform a linear approximation of the fixation probability by calculating the first derivative of the fixation probability at *δ* = 0, which we call the selection gradient. When *δ* is small, they satisfy:

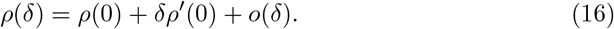

In the main text, we denote *ρ*^*′*^(0) as *µ*.

To calculate *µ*, we generalize the method proposed in [32], which is suitable for any fixed population structures. Specifically, *µ* can be expressed as:

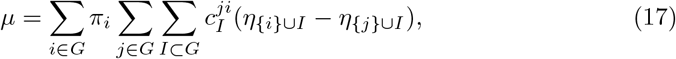

where *G* is the set of genetic sites, *π*_*i*_ represents the reproductive value of a genetic site *i*, 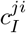 represents state *X*_*I*_ ‘s effect on selection, and *η*_*I*_ represents neutral sojourn time of state *X*_*I*_. In state *X*_*I*_, the type of the allele in site *i* is A if *i* ∈ *I*, otherwise the type of the allele is B.

Through analysis and simplifications based on structural symmetry, we found

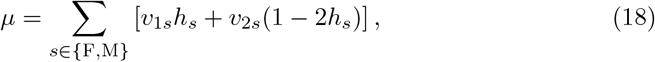

where *v*_1*s*_ measures the effect of the mutant allele at a single site of the diploid locus, while *v*_2*s*_ measures the additive effect of the two sites in the same individual.

For a detailed explanation of the above equations and a detailed derivation of the analysis, please refer to the SI Appendix, section 2.

### 4.3 Numerical simulations

To validate our analytical predictions, we ran numerical simulations, the main steps of which are described below.

1. Set the population sizes to their initial values.
2. Compute the probabilities of picking an individual whose genotype is AA, AB, or BB.
3. From the probabilities computed in step 2, compute the probabilities of picking an A or B gamete.
4. From the probabilities computed at step 3, compute the probabilities that the offspring genotype is AA, AB, or BB.
5. Generate a uniform random number between 0 and 1 and choose a new offspring proportionally to the probabilities computed in step 4.
6. Compute the probabilities of picking an individual for death whose genotype is AA, AB, or BB and having the same sex as the new offspring in the dioecious case.
7. Generate a uniform random number between 0 and 1 and choose an individual for death proportionally to the probabilities computed in step 6.
8. Update the population state by adding the new offspring and removing the dead individual.
9. Go back to step 2 until only one allele, i.e., A or B, remains in the population.

Running the numerical simulation several times allows us to calculate the number of times the mutant takes over the population and, therefore, the fixation probability.

## 5 Data availability

The authors state that all data necessary for confirming the conclusions presented in the article are represented fully within the article. Matlab implementations of numerical simulations available at https://github.com/LcMrc/MoranWithSex.

