## Appendix for "Fixation probability in a diploid sexually reproducing population"

#### 1 Description of the stochastic process model from the gene's-eye view

In a diploid population, each individual has two alleles at a genetic locus, which jointly determine the fitness of the individual. Following [1], we use the concept of genetic sites to represent arbitrary spatial locations of the alleles and their genetic structure. There are two genetic sites at a diploid locus on a single chromosome within an individual.

If an individual is chosen to reproduce, both of its alleles have equal probabilities to enter the gamete that eventually fuse with the gamete of another individual to become the new born offspring. Thus, we can consider the two alleles have the same fitness. In the gene's eye-view, alleles instead of individuals are chosen to reproduce or die. Therefore, the basic unit of selection is a genetic site. We denote the set of genetic sites in females and males as $G_{\mathrm{F}}$ and $G_{\mathrm{M}}$, respectively. The set of all genetic sites is denoted as $G = G_{\mathrm{F}} \cup G_{\mathrm{M}}$.

In a dioecious population, the probability of an allele in individual $i$ of sex $s$ to be selected is

$$b_i^s = \frac{f_i^s}{\sum_{k \in G_s} f_k^s} \qquad s \in \{\mathrm{F}, \mathrm{M}\}, i \in G_s. \tag{1}$$

In the subsequent death process, the probability of an allele in individual $j$ of sex $s$ to be selected is

$$d_j^s = \frac{1}{2N_s} \qquad s \in \{\mathrm{F}, \mathrm{M}\}. \tag{2}$$

The probability that an allele in individual $i$ replaces an allele in individual $j$ at a diploid locus is

$$e_{ij} = \frac{1}{2} b_i^s d_j^s = \frac{1}{4N_{s_j}} \frac{f_i^{s_i}}{\sum_{k \in G_{s_i}} f_k^{s_i}}. \tag{3}$$

Similarly, we can derive the expressions of the corresponding birth, death and replacement probabilities in a monoecious population,

$$b_i = \frac{2f_i}{\sum_{k \in G} f_k} \qquad i \in G, \tag{4}$$

$$d_j = \frac{1}{N}, \tag{5}$$

$$e_{ij} = \frac{1}{N} \frac{f_i}{\sum_{k \in G} f_k}. \tag{6}$$

#### 2 Ancestor random walk methods

In this subsection, we describe how to use ancestor random walk methods to calculate $\rho'(0)$, which is denoted as $\mu$. Following the modeling framework laid out in [2], we have

$$\rho'(0) = \sum_{i\in G} \pi_i \sum_{j\in G} \sum_{I\subset G} c_I^{ji} (\eta_{\{i\}\cup I} - \eta_{\{j\}\cup I}), \tag{7}$$

where $G$ is the set of genetic sites, $\pi_i$ represents the reproductive value of a genetic site $i$, $c_I^{ji}$ represents state $X_I$'s effect on selection, and $\eta_I$ represents neutral sojourn time of state $X_I$. In state $X_I$, the type of the allele in site $i$ is A if $i \in I$, otherwise the type of the allele is B. In the following we explain how to calculate each component respectively.

##### 2.1 Reproductive value

The reproductive value quantifies the expected contribution of an genetic site to the whole gene pool under neutral genetic drift, which is the unique solution of

$$\begin{cases} \sum_{j\in G} e_{ij}^{\circ} \pi_j = \sum_{j\in G} e_{ji}^{\circ} \pi_i, \\ \sum_{i\in G} \pi_i = 1, \end{cases} \tag{8}$$

where $e_{ij}^{\circ}$ is $e_{ij}$ under neutral genetic drift. In a monoecious population, $e_{ij} = \frac{1}{2N^2}$ always satisfies, so all genetic sites have the same reproductive value $\frac{1}{2N}$. In a dioecious population, the probability of an allele in individual $i$ replaces an allele in individual $j$ is $e_{ij} = \frac{1}{8N_{s_i}N_{s_j}}$. We denote the genetic sites in the set $G_{\mathrm{F}}$ and $G_{\mathrm{M}}$ as $\mathcal{F}$ and $\mathcal{M}$, respectively. Following this notation, we have

$$\begin{cases} 2N_{\mathrm{F}} \frac{1}{8N_{\mathrm{F}}N_{\mathrm{F}}} \pi_{\mathcal{F}} + 2N_{\mathrm{M}} \frac{1}{8N_{\mathrm{F}}N_{\mathrm{M}}} \pi_{\mathcal{M}} = \frac{1}{2N_{\mathrm{F}}} \pi_{\mathcal{F}}, \\ 2N_{\mathrm{F}} \pi_{\mathcal{F}} + 2N_{\mathrm{M}} \pi_{\mathcal{M}} = 1. \end{cases} \tag{9}$$

Solving equations (9) shows $\pi_{\mathcal{F}} = \pi_{\mathcal{M}} = \frac{1}{2(N_{\mathrm{F}}+N_{\mathrm{M}})}$. Namely, all genetic sites have the same reproductive value in both monoecious and dioecious populations.

##### 2.2 Effect on selection

Because $e_{ij}$ depends on the distribution of alleles in the population and their relative fitness, it is a function of $\delta$. The derivative of $e_{ij}$ with respect to $\delta$ at $\delta = 0$ is

$$e'_{ij}(\mathbf{x}) = \frac{1}{4N_{s_j}} \frac{2N_{s_i} f_i - \sum_{k\in G_{s_i}} f_k}{4N_{s_i}^2}. \tag{10}$$

We can see that $e_{ij}$ corresponds to a linear combination of all alleles. Giving the type of allele in site $i$ is $x_i$ and the type of the other allele in the same individual is $x_j$, where $x_i = 1$ if the allele in $i$ is A and $x_i = 0$ if the allele in $i$ is B. Without regard to sex, we have

$$f_i = (1+\delta)x_i x_j + (1+h\delta)x_i(1-x_j) + (1+h\delta)(1-x_i)x_j + (1-x_i)(1-x_j). \quad (11)$$

That, one's fitness only depends on two alleles. So $e'_{ij}(\mathbf{x})$ can also be represented as

$$e'_{ij}(\mathbf{x}) = \sum_{I\subset G} c_I^{ij}\mathbf{x}_I, \quad (12)$$

where $\mathbf{x}_I = \Pi_{i\in I}x_i$. As mentioned above, we only need to consider the set $|I|$ that contains either one site or two sites belonged to the same individual. In other words, $c_I^{ij} = 0$ for all $|I| > 2$.

$c_I^{ij}$ could be calculated by

$$c_I^{ij} = \sum_{J\subset I}(-1)^{|I|-|J|}e'_{ij}(\mathbf{1}_J), \quad (13)$$

where $\mathbf{1}_J$ is the state in which $x_i = 1$ if $i \in J$ and $x_i = 0$ otherwise.

Then we classify the possible $i$, $j$, and $I$ according to symmetry. In dioecious population, $i$ and $j$ can only be $\mathcal{F}$ or $\mathcal{M}$, while $I$ can be $\mathcal{F}$, $\mathcal{M}$, $\mathcal{F}\bar{\mathcal{F}}$, $\mathcal{F}\mathcal{F}'$, $\mathcal{F}\mathcal{M}$, $\mathcal{M}\bar{\mathcal{M}}$, or $\mathcal{M}\mathcal{M}'$, where $\mathcal{F}\bar{\mathcal{F}}$ ($\mathcal{M}\bar{\mathcal{M}}$) means the two sites are belonged to the same individuals, and $\mathcal{F}\mathcal{F}'$ ($\mathcal{M}\mathcal{M}'$) means they are belonged to different individuals. For convenience, we have omitted the brackets of $|I|$ here and following.

Next we list all possible cases and calculate corresponding $c_I^{ij}$. Before that, we pointed out two laws to simplify the cases that need to be listed. Firstly, the effect of $j$ on $c_I^{ij}$ is independent with $i$ and $I$, so we only need to list all possible combinations of $i$ and $I$. Secondly, if any of the element in $I$ has different sex with $i$, $c_I^{ij} = 0$ always satisfy, and we abandon these cases. In the following $\mathcal{X}$ could be $\mathcal{F}$ or $\mathcal{M}$, which sex is X.

- $i = \mathcal{X}$, $I = \mathcal{X}$ or $I = \bar{\mathcal{X}}$.

$$c_I^{ij} = \frac{1}{4N_{s_j}}\frac{2N_{\mathrm{X}}-2}{4N_{\mathrm{X}}^2}h_{\mathrm{X}}. \quad (14)$$

- $i = \mathcal{X}$, $I = \mathcal{X}'$.

$$c_I^{ij} = \frac{1}{4N_{s_j}}\frac{-2}{4N_{\mathrm{X}}^2}h_{\mathrm{X}}. \quad (15)$$

- $i = \mathcal{X}$, $I = \mathcal{X}\bar{\mathcal{X}}$.

$$c_I^{ij} = \frac{1}{4N_{s_j}}\frac{2N_{\mathrm{X}}-2}{4N_{\mathrm{X}}^2}(1-2h_{\mathrm{X}}). \quad (16)$$

- $i = \mathcal{X}$, $I = \mathcal{X}'\bar{\mathcal{X}}'$.

$$c_I^{ij} = \frac{1}{4N_{s_j}}\frac{-2}{4N_{\mathrm{X}}^2}(1-2h_{\mathrm{X}}). \quad (17)$$

In monoecious population, likewise, $i$ and $j$ can only be $\mathcal{X}$, while $I$ can be $\mathcal{X}$, $\mathcal{X}\bar{\mathcal{X}}$, or $\mathcal{X}\mathcal{X}'$. To calculate $c_I^{ij}$, we only need replace $4N_{s_j}$ and $N_{\mathrm{X}}$ with $N$.

#### 2.3 Neutral sojourn time

Neutral sojourn time measures the mean number of steps in which all sites in $I$ have type $A$ prior to absorption, which is the unique solution of

$$\begin{cases} \eta_I = E^\circ[\sum_{i\in I}\pi_i - \xi_I] + \sum_\alpha p^\circ_\alpha \eta_{\alpha I}, \\ \sum_{i\in G}\pi_i\eta_i = 0, \end{cases} \tag{18}$$

where $E^\circ$ is the expectation under a certain mutant-appearance distribution, and $p^\circ_\alpha$ is the probability that $I$ change into $\alpha(I)$ in a single step under neutral selection, with the mapping $\alpha$.

The situation we consider is mutation happens in female, in which we can get $E^\circ[\pi_{\mathcal{F}} - \xi_{\mathcal{F}}] = \frac{1}{2(N_{\mathrm{F}}+N_{\mathrm{M}})} - \frac{1}{2N_{\mathrm{F}}}$, and $E^\circ[\pi_{\mathcal{M}} - \xi_{\mathcal{M}}] = \frac{1}{2(N_{\mathrm{F}}+N_{\mathrm{M}})}$. Besides, if $|I|>1$, $\xi_I = 0$ always satisfy, so $E^\circ[\sum_{i\in I}\pi_i - \xi_I] = \frac{1}{2(N_{\mathrm{F}}+N_{\mathrm{M}})}$. Because $c_I^{ij} = 0$ for all $|I|>2$, here we only need to solve $\eta_I$ with $|I| \le 3$. Considering symmetry again, all sets that we should consider include $\mathcal{F}$, $\mathcal{M}$, $\mathcal{F}\bar{\mathcal{F}}$, $\mathcal{F}\mathcal{F}'$, $\mathcal{M}\bar{\mathcal{M}}$, $\mathcal{M}\mathcal{M}'$, $\mathcal{F}\mathcal{M}$, $\mathcal{F}\bar{\mathcal{F}}\mathcal{F}'$, $\mathcal{F}\mathcal{F}'\mathcal{F}''$, $\mathcal{F}\bar{\mathcal{F}}\mathcal{M}$, $\mathcal{F}\mathcal{F}'\mathcal{M}$, $\mathcal{M}\bar{\mathcal{M}}\mathcal{M}'$, $\mathcal{M}\mathcal{M}'\mathcal{M}''$, $\mathcal{M}\bar{\mathcal{M}}\mathcal{F}$, and $\mathcal{M}\mathcal{M}'\mathcal{F}$.

$$\left\{\begin{array}{l}
2N_{\mathrm{F}}\eta_{\mathcal{F}} + 2N_{\mathrm{M}}\eta_{\mathcal{M}} = 0, \\
\frac{1}{2N_{\mathrm{F}}}\eta_{\mathcal{F}} = \frac{1}{2(N_{\mathrm{F}}+N_{\mathrm{M}})} - \frac{1}{2N_{\mathrm{F}}} + \frac{1}{4N_{\mathrm{F}}}\eta_{\mathcal{F}} + \frac{1}{4N_{\mathrm{F}}}\eta_{\mathcal{M}}, \\
\frac{1}{N_{\mathrm{F}}}\eta_{\mathcal{F}\bar{\mathcal{F}}} = \frac{1}{2(N_{\mathrm{F}}+N_{\mathrm{M}})} + \frac{1}{2N_{\mathrm{F}}}\eta_{\mathcal{F}\mathcal{M}}, \\
\frac{1}{2N_{\mathrm{M}}}\eta_{\mathcal{M}\bar{\mathcal{M}}} = \frac{1}{2(N_{\mathrm{F}}+N_{\mathrm{M}})} + \frac{1}{2N_{\mathrm{M}}}\eta_{\mathcal{F}\mathcal{M}}, \\
\frac{1}{2N_{\mathrm{F}}}\eta_{\mathcal{F}\mathcal{F}'} = \frac{1}{2(N_{\mathrm{F}}+N_{\mathrm{M}})} + \frac{1}{4N_{\mathrm{F}}^2}\eta_{\mathcal{F}} + \frac{1}{4N_{\mathrm{F}}^2}\eta_{\mathcal{F}\bar{\mathcal{F}}} + \frac{N_{\mathrm{F}}-1}{2N_{\mathrm{F}}^2}\eta_{\mathcal{F}\mathcal{F}'} + \frac{1}{2N_{\mathrm{F}}}\eta_{\mathcal{F}\mathcal{M}}, \\
\frac{1}{N_{\mathrm{M}}}\eta_{\mathcal{M}\mathcal{M}'} = \frac{1}{2(N_{\mathrm{F}}+N_{\mathrm{M}})} + \frac{1}{4N_{\mathrm{M}}^2}\eta_{\mathcal{M}} + \frac{1}{4N_{\mathrm{M}}^2}\eta_{\mathcal{M}\bar{\mathcal{M}}} + \frac{N_{\mathrm{M}}-1}{2N_{\mathrm{M}}^2}\eta_{\mathcal{M}\mathcal{M}'} + \frac{1}{2N_{\mathrm{M}}}\eta_{\mathcal{F}\mathcal{M}}, \\
(\frac{1}{2N_{\mathrm{F}}} + \frac{1}{2N_{\mathrm{M}}})\eta_{\mathcal{F}\mathcal{M}} = \frac{1}{2(N_{\mathrm{F}}+N_{\mathrm{M}})} + (\frac{1}{4N_{\mathrm{F}}} + \frac{1}{4N_{\mathrm{M}}})\eta_{\mathcal{F}\mathcal{M}} + \frac{1}{8N_{\mathrm{F}}N_{\mathrm{M}}}\eta_{\mathcal{M}} + \frac{1}{8N_{\mathrm{F}}N_{\mathrm{M}}}\eta_{\mathcal{M}\bar{\mathcal{M}}} + \frac{N_{\mathrm{M}}-1}{4N_{\mathrm{F}}N_{\mathrm{M}}}\eta_{\mathcal{M}\mathcal{M}'} \\
\qquad + \frac{1}{8N_{\mathrm{F}}N_{\mathrm{M}}}\eta_{\mathcal{F}} + \frac{1}{8N_{\mathrm{F}}N_{\mathrm{M}}}\eta_{\mathcal{F}\bar{\mathcal{F}}} + \frac{N_{\mathrm{F}}-1}{4N_{\mathrm{F}}N_{\mathrm{M}}}\eta_{\mathcal{F}\mathcal{F}'}, \\
\frac{1}{N_{\mathrm{F}}}\eta_{\mathcal{F}\bar{\mathcal{F}}\mathcal{F}'} = \frac{1}{2(N_{\mathrm{F}}+N_{\mathrm{M}})} + \frac{1}{4N_{\mathrm{F}}^2}\eta_{\mathcal{F}\mathcal{M}} + \frac{1}{4N_{\mathrm{F}}^2}\eta_{\mathcal{F}\bar{\mathcal{F}}\mathcal{M}} + (\frac{N_{\mathrm{F}}-1}{2N_{\mathrm{F}}^2} + \frac{1}{4N_{\mathrm{F}}^2})\eta_{\mathcal{F}\mathcal{F}'\mathcal{M}} + \frac{1}{4N_{\mathrm{F}}^2}\eta_{\mathcal{F}\bar{\mathcal{F}}} + \frac{N_{\mathrm{F}}-1}{4N_{\mathrm{F}}^2}\eta_{\mathcal{F}\bar{\mathcal{F}}\mathcal{F}'}, \\
\frac{1}{N_{\mathrm{M}}}\eta_{\mathcal{M}\bar{\mathcal{M}}\mathcal{M}'} = \frac{1}{2(N_{\mathrm{F}}+N_{\mathrm{M}})} + \frac{1}{4N_{\mathrm{M}}^2}\eta_{\mathcal{F}\mathcal{M}} + \frac{1}{4N_{\mathrm{M}}^2}\eta_{\mathcal{M}\bar{\mathcal{M}}\mathcal{F}} + (\frac{N_{\mathrm{M}}-1}{2N_{\mathrm{M}}^2} + \frac{1}{4N_{\mathrm{M}}^2})\eta_{\mathcal{M}\mathcal{M}'\mathcal{F}} \\
\qquad + \frac{1}{4N_{\mathrm{M}}^2}\eta_{\mathcal{M}\bar{\mathcal{M}}} + \frac{N_{\mathrm{M}}-1}{4N_{\mathrm{M}}^2}\eta_{\mathcal{M}\bar{\mathcal{M}}\mathcal{M}'}, \\
\frac{3}{2N_{\mathrm{F}}}\eta_{\mathcal{F}\mathcal{F}'\mathcal{F}''} = \frac{1}{2(N_{\mathrm{F}}+N_{\mathrm{M}})} + \frac{3}{4N_{\mathrm{F}}^2}\eta_{\mathcal{F}\mathcal{F}'} + \frac{3}{4N_{\mathrm{F}}^2}\eta_{\mathcal{F}\bar{\mathcal{F}}\mathcal{F}'} + \frac{3N_{\mathrm{F}}-6}{4N_{\mathrm{F}}^2}\eta_{\mathcal{F}\mathcal{F}'\mathcal{F}''} + \frac{3}{4N_{\mathrm{F}}}\eta_{\mathcal{F}\mathcal{F}'\mathcal{M}}, \\
\frac{3}{2N_{\mathrm{M}}}\eta_{\mathcal{M}\mathcal{M}'\mathcal{M}''} = \frac{1}{2(N_{\mathrm{F}}+N_{\mathrm{M}})} + \frac{3}{4N_{\mathrm{M}}^2}\eta_{\mathcal{M}\mathcal{M}'} + \frac{3}{4N_{\mathrm{M}}^2}\eta_{\mathcal{M}\bar{\mathcal{M}}\mathcal{M}'} + \frac{3N_{\mathrm{M}}-6}{4N_{\mathrm{M}}^2}\eta_{\mathcal{M}\mathcal{M}'\mathcal{M}''} + \frac{3}{4N_{\mathrm{M}}}\eta_{\mathcal{M}\mathcal{M}'\mathcal{F}}, \\
(\frac{1}{2N_{\mathrm{F}}} + \frac{1}{2N_{\mathrm{M}}})\eta_{\mathcal{F}\bar{\mathcal{F}}\mathcal{M}} = \frac{1}{2(N_{\mathrm{F}}+N_{\mathrm{M}})} + \frac{1}{4N_{\mathrm{F}}N_{\mathrm{M}}}\eta_{\mathcal{F}\mathcal{M}} + \frac{1}{4N_{\mathrm{F}}N_{\mathrm{M}}}\eta_{\mathcal{M}\bar{\mathcal{M}}\mathcal{F}} + \frac{N_{\mathrm{M}}-1}{2N_{\mathrm{F}}N_{\mathrm{M}}}\eta_{\mathcal{M}\mathcal{M}'\mathcal{F}} + \frac{1}{4N_{\mathrm{F}}N_{\mathrm{M}}}\eta_{\mathcal{F}\bar{\mathcal{F}}} \\
\qquad + \frac{N_{\mathrm{F}}-1}{4N_{\mathrm{F}}N_{\mathrm{M}}}\eta_{\mathcal{F}\bar{\mathcal{F}}\mathcal{F}'} + \frac{1}{4N_{\mathrm{M}}}\eta_{\mathcal{F}\bar{\mathcal{F}}\mathcal{M}}, \\
(\frac{1}{2N_{\mathrm{F}}} + \frac{1}{2N_{\mathrm{M}}})\eta_{\mathcal{M}\bar{\mathcal{M}}\mathcal{F}} = \frac{1}{2(N_{\mathrm{F}}+N_{\mathrm{M}})} + \frac{1}{4N_{\mathrm{F}}N_{\mathrm{M}}}\eta_{\mathcal{F}\mathcal{M}} + \frac{1}{4N_{\mathrm{F}}N_{\mathrm{M}}}\eta_{\mathcal{F}\bar{\mathcal{F}}\mathcal{M}} \\
\qquad + \frac{N_{\mathrm{F}}-1}{2N_{\mathrm{F}}N_{\mathrm{M}}}\eta_{\mathcal{F}\mathcal{F}'\mathcal{M}} + \frac{1}{4N_{\mathrm{F}}N_{\mathrm{M}}}\eta_{\mathcal{M}\bar{\mathcal{M}}} + \frac{N_{\mathrm{M}}-1}{4N_{\mathrm{F}}N_{\mathrm{M}}}\eta_{\mathcal{M}\bar{\mathcal{M}}\mathcal{M}'} + \frac{1}{4N_{\mathrm{F}}}\eta_{\mathcal{M}\bar{\mathcal{M}}\mathcal{F}}, \\
(\frac{1}{N_{\mathrm{F}}} + \frac{1}{2N_{\mathrm{M}}})\eta_{\mathcal{F}\mathcal{F}'\mathcal{M}} = \frac{1}{2(N_{\mathrm{F}}+N_{\mathrm{M}})} + (\frac{1}{4N_{\mathrm{F}}^2} + \frac{1}{4N_{\mathrm{F}}N_{\mathrm{M}}})\eta_{\mathcal{F}\mathcal{M}} + \frac{1}{4N_{\mathrm{F}}^2}\eta_{\mathcal{F}\bar{\mathcal{F}}\mathcal{M}} + (\frac{N_{\mathrm{F}}-1}{2N_{\mathrm{F}}^2} + \frac{1}{4N_{\mathrm{M}}})\eta_{\mathcal{F}\mathcal{F}'\mathcal{M}} \\
\qquad + \frac{1}{4N_{\mathrm{F}}N_{\mathrm{M}}}\eta_{\mathcal{M}\bar{\mathcal{M}}\mathcal{F}} + \frac{N_{\mathrm{M}}-1}{2N_{\mathrm{F}}N_{\mathrm{M}}}\eta_{\mathcal{M}\mathcal{M}'\mathcal{F}} + \frac{1}{4N_{\mathrm{F}}N_{\mathrm{M}}}\eta_{\mathcal{F}\mathcal{F}'} + \frac{1}{4N_{\mathrm{F}}N_{\mathrm{M}}}\eta_{\mathcal{F}\bar{\mathcal{F}}\mathcal{F}'} + \frac{N_{\mathrm{F}}-2}{4N_{\mathrm{F}}N_{\mathrm{M}}}\eta_{\mathcal{F}\mathcal{F}'\mathcal{F}''}, \\
(\frac{1}{N_{\mathrm{M}}} + \frac{1}{2N_{\mathrm{F}}})\eta_{\mathcal{M}\mathcal{M}'\mathcal{F}} = \frac{1}{2(N_{\mathrm{F}}+N_{\mathrm{M}})} + (\frac{1}{4N_{\mathrm{M}}^2} + \frac{1}{4N_{\mathrm{F}}N_{\mathrm{M}}})\eta_{\mathcal{F}\mathcal{M}} + \frac{1}{4N_{\mathrm{M}}^2}\eta_{\mathcal{M}\bar{\mathcal{M}}\mathcal{F}} + (\frac{N_{\mathrm{M}}-1}{2N_{\mathrm{M}}^2} + \frac{1}{4N_{\mathrm{F}}})\eta_{\mathcal{M}\mathcal{M}'\mathcal{F}} \\
\qquad + \frac{1}{4N_{\mathrm{F}}N_{\mathrm{M}}}\eta_{\mathcal{F}\bar{\mathcal{F}}\mathcal{M}} + \frac{N_{\mathrm{F}}-1}{2N_{\mathrm{F}}N_{\mathrm{M}}}\eta_{\mathcal{F}\mathcal{F}'\mathcal{M}} + \frac{1}{4N_{\mathrm{F}}N_{\mathrm{M}}}\eta_{\mathcal{M}\mathcal{M}'} + \frac{1}{4N_{\mathrm{F}}N_{\mathrm{M}}}\eta_{\mathcal{M}\bar{\mathcal{M}}\mathcal{M}'} + \frac{N_{\mathrm{M}}-2}{4N_{\mathrm{F}}N_{\mathrm{M}}}\eta_{\mathcal{M}\mathcal{M}'\mathcal{M}''}.
\end{array}\right. \tag{19}$$

We use maple to solve the above equations, and we won't show the exactly solution here because the specific form of the solution is too complicated.

In monoecious population, since each site in the population is homogeneous, $E^{\circ}$ is always $\frac{1}{2N}$, so $\eta_i$ is always 0. For $|I|>1$, we only need to replace $\mathcal{F}$ and $\mathcal{M}$ in the above equations with $\mathcal{X}$ (Half of the equations are repetitive).

#### 2.4 Detailed expression

We have calculated all possible $\pi_i$, $c_I^{ji}$, $\eta_{i\cup I}$, and $\eta_{j\cup I}$. Substitute all $i$,$j$,$I$ into the expression of $\rho'(0)$, we can get its analytical form:

$$\begin{aligned}\mu =&\frac{N_{\mathrm{F}}^2N_{\mathrm{M}}+N_{\mathrm{F}}N_{\mathrm{M}}^2+N_{\mathrm{F}}^2+N_{\mathrm{F}}N_{\mathrm{M}}-N_{\mathrm{M}}^2-N_{\mathrm{F}}-2N_{\mathrm{M}}}{4N_{\mathrm{F}}^2N_{\mathrm{M}}+4N_{\mathrm{F}}N_{\mathrm{M}}^2+2N_{\mathrm{F}}^2+4N_{\mathrm{F}}N_{\mathrm{M}}+2N_{\mathrm{M}}^2}h_F+\\ &\frac{N_{\mathrm{F}}^2N_{\mathrm{M}}+N_{\mathrm{F}}N_{\mathrm{M}}^2-N_{\mathrm{F}}^2-N_{\mathrm{F}}N_{\mathrm{M}}+N_{\mathrm{M}}^2-N_{\mathrm{M}}}{4N_{\mathrm{F}}^2N_{\mathrm{M}}+4N_{\mathrm{F}}N_{\mathrm{M}}^2+2N_{\mathrm{F}}^2+4N_{\mathrm{F}}N_{\mathrm{M}}+2N_{\mathrm{M}}^2}h_M+\\ &\frac{num_F}{den_F}(1-2h_F)+\frac{num_M}{den_M}(1-2h_M),\end{aligned} \tag{20}$$

where $num_F = 12N_{\mathrm{F}}^9N_{\mathrm{M}}^3+76N_{\mathrm{F}}^8N_{\mathrm{M}}^4+168N_{\mathrm{F}}^7N_{\mathrm{M}}^5+168N_{\mathrm{F}}^6N_{\mathrm{M}}^6+76N_{\mathrm{F}}^5N_{\mathrm{M}}^7+12N_{\mathrm{F}}^4N_{\mathrm{M}}^8+40N_{\mathrm{F}}^9N_{\mathrm{M}}^2+280N_{\mathrm{F}}^8N_{\mathrm{M}}^3+650N_{\mathrm{F}}^7N_{\mathrm{M}}^4+632N_{\mathrm{F}}^6N_{\mathrm{M}}^5+340N_{\mathrm{F}}^5N_{\mathrm{M}}^6+96N_{\mathrm{F}}^4N_{\mathrm{M}}^7+10N_{\mathrm{F}}^3N_{\mathrm{M}}^8+36N_{\mathrm{F}}^9N_{\mathrm{M}}+324N_{\mathrm{F}}^8N_{\mathrm{M}}^2+847N_{\mathrm{F}}^7N_{\mathrm{M}}^3+712N_{\mathrm{F}}^6N_{\mathrm{M}}^4+190N_{\mathrm{F}}^5N_{\mathrm{M}}^5-152N_{\mathrm{F}}^4N_{\mathrm{M}}^6-137N_{\mathrm{F}}^3N_{\mathrm{M}}^7-28N_{\mathrm{F}}^2N_{\mathrm{M}}^8+8N_{\mathrm{F}}^9+112N_{\mathrm{F}}^8N_{\mathrm{M}}+314N_{\mathrm{F}}^7N_{\mathrm{M}}^2+16N_{\mathrm{F}}^6N_{\mathrm{M}}^3-482N_{\mathrm{F}}^5N_{\mathrm{M}}^4-628N_{\mathrm{F}}^4N_{\mathrm{M}}^5-410N_{\mathrm{F}}^3N_{\mathrm{M}}^6-84N_{\mathrm{F}}^2N_{\mathrm{M}}^7+2N_{\mathrm{F}}N_{\mathrm{M}}^8+8N_{\mathrm{F}}^8-5N_{\mathrm{F}}^7N_{\mathrm{M}}-264N_{\mathrm{F}}^6N_{\mathrm{M}}^2-549N_{\mathrm{F}}^5N_{\mathrm{M}}^3-714N_{\mathrm{F}}^4N_{\mathrm{M}}^4-461N_{\mathrm{F}}^3N_{\mathrm{M}}^5-26N_{\mathrm{F}}^2N_{\mathrm{M}}^6+39N_{\mathrm{F}}N_{\mathrm{M}}^7+4N_{\mathrm{M}}^8-6N_{\mathrm{F}}^7-72N_{\mathrm{F}}^6N_{\mathrm{M}}-202N_{\mathrm{F}}^5N_{\mathrm{M}}^2-444N_{\mathrm{F}}^4N_{\mathrm{M}}^3-264N_{\mathrm{F}}^3N_{\mathrm{M}}^4+58N_{\mathrm{F}}^2N_{\mathrm{M}}^5+72N_{\mathrm{F}}N_{\mathrm{M}}^6+10N_{\mathrm{M}}^7-8N_{\mathrm{F}}^6-55N_{\mathrm{F}}^5N_{\mathrm{M}}-186N_{\mathrm{F}}^4N_{\mathrm{M}}^2-152N_{\mathrm{F}}^3N_{\mathrm{M}}^3+18N_{\mathrm{F}}^2N_{\mathrm{M}}^4+39N_{\mathrm{F}}N_{\mathrm{M}}^5+8N_{\mathrm{M}}^6-2N_{\mathrm{F}}^5-16N_{\mathrm{F}}^4N_{\mathrm{M}}-26N_{\mathrm{F}}^3N_{\mathrm{M}}^2-10N_{\mathrm{F}}^2N_{\mathrm{M}}^3+4N_{\mathrm{F}}N_{\mathrm{M}}^4+2N_{\mathrm{M}}^5$,

$num_M = 12N_{\mathrm{F}}^8N_{\mathrm{M}}^4+76N_{\mathrm{F}}^7N_{\mathrm{M}}^5+168N_{\mathrm{F}}^6N_{\mathrm{M}}^6+168N_{\mathrm{F}}^5N_{\mathrm{M}}^7+76N_{\mathrm{F}}^4N_{\mathrm{M}}^8+12N_{\mathrm{F}}^3N_{\mathrm{M}}^9+10N_{\mathrm{F}}^8N_{\mathrm{M}}^3+96N_{\mathrm{F}}^7N_{\mathrm{M}}^4+340N_{\mathrm{F}}^6N_{\mathrm{M}}^5+632N_{\mathrm{F}}^5N_{\mathrm{M}}^6+650N_{\mathrm{F}}^4N_{\mathrm{M}}^7+280N_{\mathrm{F}}^3N_{\mathrm{M}}^8+40N_{\mathrm{F}}^2N_{\mathrm{M}}^9-28N_{\mathrm{F}}^8N_{\mathrm{M}}^2-137N_{\mathrm{F}}^7N_{\mathrm{M}}^3-152N_{\mathrm{F}}^6N_{\mathrm{M}}^4+190N_{\mathrm{F}}^5N_{\mathrm{M}}^5+712N_{\mathrm{F}}^4N_{\mathrm{M}}^6+847N_{\mathrm{F}}^3N_{\mathrm{M}}^7+324N_{\mathrm{F}}^2N_{\mathrm{M}}^8+36N_{\mathrm{F}}N_{\mathrm{M}}^9+2N_{\mathrm{F}}^8N_{\mathrm{M}}-84N_{\mathrm{F}}^7N_{\mathrm{M}}^2-446N_{\mathrm{F}}^6N_{\mathrm{M}}^3-712N_{\mathrm{F}}^5N_{\mathrm{M}}^4-326N_{\mathrm{F}}^4N_{\mathrm{M}}^5+292N_{\mathrm{F}}^3N_{\mathrm{M}}^6+386N_{\mathrm{F}}^2N_{\mathrm{M}}^7+112N_{\mathrm{F}}N_{\mathrm{M}}^8+8N_{\mathrm{M}}^9+4N_{\mathrm{F}}^8+39N_{\mathrm{F}}^7N_{\mathrm{M}}+22N_{\mathrm{F}}^6N_{\mathrm{M}}^2-389N_{\mathrm{F}}^5N_{\mathrm{M}}^3-954N_{\mathrm{F}}^4N_{\mathrm{M}}^4-789N_{\mathrm{F}}^3N_{\mathrm{M}}^5-216N_{\mathrm{F}}^2N_{\mathrm{M}}^6+19N_{\mathrm{F}}N_{\mathrm{M}}^7+8N_{\mathrm{M}}^8+10N_{\mathrm{F}}^7+60N_{\mathrm{F}}^6N_{\mathrm{M}}+70N_{\mathrm{F}}^5N_{\mathrm{M}}^2-240N_{\mathrm{F}}^4N_{\mathrm{M}}^3-660N_{\mathrm{F}}^3N_{\mathrm{M}}^4-454N_{\mathrm{F}}^2N_{\mathrm{M}}^5-108N_{\mathrm{F}}N_{\mathrm{M}}^6-6N_{\mathrm{M}}^7+8N_{\mathrm{F}}^6+39N_{\mathrm{F}}^5N_{\mathrm{M}}+66N_{\mathrm{F}}^4N_{\mathrm{M}}^2-8N_{\mathrm{F}}^3N_{\mathrm{M}}^3-90N_{\mathrm{F}}^2N_{\mathrm{M}}^4-55N_{\mathrm{F}}N_{\mathrm{M}}^5-8N_{\mathrm{M}}^6+2N_{\mathrm{F}}^5+16N_{\mathrm{F}}^4N_{\mathrm{M}}+26N_{\mathrm{F}}^3N_{\mathrm{M}}^2+10N_{\mathrm{F}}^2N_{\mathrm{M}}^3-4N_{\mathrm{F}}N_{\mathrm{M}}^4-2N_{\mathrm{M}}^5$,

$den_F = 144N_{\mathrm{F}}^9N_{\mathrm{M}}^3+912N_{\mathrm{F}}^8N_{\mathrm{M}}^4+2016N_{\mathrm{F}}^7N_{\mathrm{M}}^5+2016N_{\mathrm{F}}^6N_{\mathrm{M}}^6+912N_{\mathrm{F}}^5N_{\mathrm{M}}^7+144N_{\mathrm{F}}^4N_{\mathrm{M}}^8+408N_{\mathrm{F}}^9N_{\mathrm{M}}^2+2976N_{\mathrm{F}}^8N_{\mathrm{M}}^3+8040N_{\mathrm{F}}^7N_{\mathrm{M}}^4+10944N_{\mathrm{F}}^6N_{\mathrm{M}}^5+8040N_{\mathrm{F}}^5N_{\mathrm{M}}^6+2976N_{\mathrm{F}}^4N_{\mathrm{M}}^7+408N_{\mathrm{F}}^3N_{\mathrm{M}}^8+264N_{\mathrm{F}}^9N_{\mathrm{M}}+2832N_{\mathrm{F}}^8N_{\mathrm{M}}^2+10560N_{\mathrm{F}}^7N_{\mathrm{M}}^3+19368N_{\mathrm{F}}^6N_{\mathrm{M}}^4+19368N_{\mathrm{F}}^5N_{\mathrm{M}}^5+10560N_{\mathrm{F}}^4N_{\mathrm{M}}^6+2832N_{\mathrm{F}}^3N_{\mathrm{M}}^7+264N_{\mathrm{F}}^2N_{\mathrm{M}}^8+48N_{\mathrm{F}}^9+960N_{\mathrm{F}}^8N_{\mathrm{M}}+5328N_{\mathrm{F}}^7N_{\mathrm{M}}^2+14016N_{\mathrm{F}}^6N_{\mathrm{M}}^3+19200N_{\mathrm{F}}^5N_{\mathrm{M}}^4+14016N_{\mathrm{F}}^4N_{\mathrm{M}}^5+5328N_{\mathrm{F}}^3N_{\mathrm{M}}^6+960N_{\mathrm{F}}^2N_{\mathrm{M}}^7+48N_{\mathrm{F}}N_{\mathrm{M}}^8+96N_{\mathrm{F}}^8+984N_{\mathrm{F}}^7N_{\mathrm{M}}+4056N_{\mathrm{F}}^6N_{\mathrm{M}}^2+7920N_{\mathrm{F}}^5N_{\mathrm{M}}^3+7920N_{\mathrm{F}}^4N_{\mathrm{M}}^4+4056N_{\mathrm{F}}^3N_{\mathrm{M}}^5+984N_{\mathrm{F}}^2N_{\mathrm{M}}^6+96N_{\mathrm{F}}N_{\mathrm{M}}^7+48N_{\mathrm{F}}^7+384N_{\mathrm{F}}^6N_{\mathrm{M}}+1104N_{\mathrm{F}}^5N_{\mathrm{M}}^2+1536N_{\mathrm{F}}^4N_{\mathrm{M}}^3+1104N_{\mathrm{F}}^3N_{\mathrm{M}}^4+384N_{\mathrm{F}}^2N_{\mathrm{M}}^5+48N_{\mathrm{F}}N_{\mathrm{M}}^6$,

$den_M = 144N_{\mathrm{F}}^8N_{\mathrm{M}}^4+912N_{\mathrm{F}}^7N_{\mathrm{M}}^5+2016N_{\mathrm{F}}^6N_{\mathrm{M}}^6+2016N_{\mathrm{F}}^5N_{\mathrm{M}}^7+912N_{\mathrm{F}}^4N_{\mathrm{M}}^8+144N_{\mathrm{F}}^3N_{\mathrm{M}}^9+408N_{\mathrm{F}}^8N_{\mathrm{M}}^3+2976N_{\mathrm{F}}^7N_{\mathrm{M}}^4+8040N_{\mathrm{F}}^6N_{\mathrm{M}}^5+10944N_{\mathrm{F}}^5N_{\mathrm{M}}^6+8040N_{\mathrm{F}}^4N_{\mathrm{M}}^7+2976N_{\mathrm{F}}^3N_{\mathrm{M}}^8+408N_{\mathrm{F}}^2N_{\mathrm{M}}^9+264N_{\mathrm{F}}^8N_{\mathrm{M}}^2+2832N_{\mathrm{F}}^7N_{\mathrm{M}}^3+10560N_{\mathrm{F}}^6N_{\mathrm{M}}^4+19368N_{\mathrm{F}}^5N_{\mathrm{M}}^5+19368N_{\mathrm{F}}^4N_{\mathrm{M}}^6+10560N_{\mathrm{F}}^3N_{\mathrm{M}}^7+2832N_{\mathrm{F}}^2N_{\mathrm{M}}^8+264N_{\mathrm{F}}N_{\mathrm{M}}^9+48N_{\mathrm{F}}^8N_{\mathrm{M}}+960N_{\mathrm{F}}^7N_{\mathrm{M}}^2+5328N_{\mathrm{F}}^6N_{\mathrm{M}}^3+14016N_{\mathrm{F}}^5N_{\mathrm{M}}^4+19200N_{\mathrm{F}}^4N_{\mathrm{M}}^5+14016N_{\mathrm{F}}^3N_{\mathrm{M}}^6+5328N_{\mathrm{F}}^2N_{\mathrm{M}}^7+960N_{\mathrm{F}}N_{\mathrm{M}}^8+$

$48N_{\mathrm{M}}^9+96N_{\mathrm{F}}^7N_{\mathrm{M}}+984N_{\mathrm{F}}^6N_{\mathrm{M}}^2+4056N_{\mathrm{F}}^5N_{\mathrm{M}}^3+7920N_{\mathrm{F}}^4N_{\mathrm{M}}^4+7920N_{\mathrm{F}}^3N_{\mathrm{M}}^5+4056N_{\mathrm{F}}^2N_{\mathrm{M}}^6+984N_{\mathrm{F}}N_{\mathrm{M}}^7+96N_{\mathrm{M}}^8+48N_{\mathrm{F}}^6N_{\mathrm{M}}+384N_{\mathrm{F}}^5N_{\mathrm{M}}^2+1104N_{\mathrm{F}}^4N_{\mathrm{M}}^3+1536N_{\mathrm{F}}^3N_{\mathrm{M}}^4+1104N_{\mathrm{F}}^2N_{\mathrm{M}}^5+384N_{\mathrm{F}}N_{\mathrm{M}}^6+48N_{\mathrm{M}}^7$.

For convenience, the above formula is abbreviated as

$$\mu = v_{1\mathrm{F}}h_{\mathrm{F}} + v_{1\mathrm{M}}h_{\mathrm{M}} + v_{2\mathrm{F}}(1-2h_{\mathrm{F}}) + v_{2\mathrm{M}}(1-2h_{\mathrm{M}}). \tag{21}$$

### 3 Further explanation of the main results

#### 3.1 Comparison between haploid and diploid populations

We suppose that the ratio of B to A is $U$. In haploid population, the probability that $A$ is chosen to give birth is

$$\beta_{\mathrm{A}}^{H}(\delta) = \frac{1+\delta}{1+\delta+U}. \tag{22}$$

When $\delta \to 0$, through Taylor expansion, we have

$$\lim_{\delta\to 0}\beta_{\mathrm{A}}^{H}(\delta) = \frac{1}{1+U} + \delta\frac{U}{(1+U)^2} + o(\delta). \tag{23}$$

In diploid population, when $N$ is large enough, the fractions of three phenotypes satisfy Hardy-Weinberg equilibrium [3], that is $r_{\mathrm{AA}} : r_{\mathrm{AB}} : r_{\mathrm{BB}} = 1 : 2U : U^2$, and the probability that $A$ is chosen to give birth is,

$$\beta_{\mathrm{A}}^{D}(\delta) = \frac{1+\delta+U(1+h\delta)}{1+\delta+2U(1+h\delta)+U^2}. \tag{24}$$

When $\delta \to 0$, through Taylor expansion, we have

$$\lim_{\delta\to 0}\beta_{\mathrm{A}}^{D}(\delta) = \frac{1}{1+U} + \delta\frac{hU^2+(1-h)U}{(1+U)^3} + o(\delta). \tag{25}$$

According to Ohtsuki et al.'s work [4], the average values of $U$ during the whole evolutionary process is 2 when $\delta \to 0$, substituting $U = 2$ into equation (25), we have

$$\lim_{\delta\to 0}\overline{\beta}_{\mathrm{A}}^{H}(\delta) = \frac{1}{3} + \frac{2}{9}\delta + o(\delta), \tag{26}$$

and

$$\lim_{\delta\to 0}\overline{\beta}_{\mathrm{A}}^{D}(\delta) = \frac{1}{3} + \frac{2h+2}{27}\delta + o(\delta). \tag{27}$$

When $h = 2$, we have $\lim_{\delta\to 0}\overline{\beta}_{\mathrm{A}}^{DD}(\delta) = \lim_{\delta\to 0}\overline{\beta}_{\mathrm{A}}^{H}(\delta)$, and when $h = -1$, we have $\lim_{\delta\to 0}\overline{\beta}_{\mathrm{A}}^{DD}(\delta) = \overline{\beta}_{\mathrm{A}}^{H}(0)$, which are consistent with the main results.

#### 3.2 Effect of biased sex ratio

We take $h = 1/2$ as an example. When $h = 1/2$ we have

$$\mu = \frac{2N_\mathrm{F}^2 N_\mathrm{M} + 2N_\mathrm{F} N_\mathrm{M}^2 - N_\mathrm{F} - 3N_\mathrm{M}}{4(N_\mathrm{F} + N_\mathrm{M})(2N_\mathrm{F} N_\mathrm{M} + N_\mathrm{F} + N_\mathrm{M})}. \tag{28}$$

We assume that $N_\mathrm{F} + N_\mathrm{M} = 2N$, thus $\mu$ could be rewritten as a function of $N_\mathrm{F}$

$$\begin{aligned} \mu &= \frac{4N_\mathrm{F}(2N - N_\mathrm{F})N - 6N + 2N_\mathrm{F}}{8N(2N_\mathrm{F}(2N - N_\mathrm{F}) + 2N)} \\ &= \frac{-4NN_\mathrm{F}^2 + (8N^2 + 2)N_\mathrm{F} - 6N}{-16NN_\mathrm{F}^2 + 32N^2 N_\mathrm{F} + 16N^2}. \end{aligned} \tag{29}$$

By calculating the derivative on $N_\mathrm{F}$, it is found that the maximum point of $\mu$ is $N_\mathrm{F}^* = 2N^2 + 3N - \sqrt{4N^4 + 8N^3 + 3N^2 - N}$. When $N_\mathrm{F} < N_\mathrm{F}^*$, $\mu$ increases monotonically with $N_\mathrm{F}$. When $N_\mathrm{F} > N_\mathrm{F}^*$, $\mu$ decreases monotonically with $N_\mathrm{F}$. When $N$ is large (but finite), $N_\mathrm{F}^* \approx N$, that is, biased sex ratios tend to suppress natural selection.

#### 3.3 Effects of sex-specific dominance coefficients

We take $h_\mathrm{F} = 1$, $h_\mathrm{M} = 0$ and $h_\mathrm{F} = 0$, $h_\mathrm{M} = 1$ as two examples. We assume that $N_\mathrm{F} + N_\mathrm{M} = 2N$ and calculating the derivative on $N_\mathrm{F}$, getting that $v'_{1\mathrm{M}} > 0$, $v'_{2\mathrm{M}} > 0$, $v'_{1\mathrm{F}} < 0$, and $v'_{2\mathrm{F}} < 0$.

If we let $h_\mathrm{F} = 1$ and $h_\mathrm{M} = 0$, we have

$$\mu' = v'_{1_\mathrm{F}} - v'_{2_\mathrm{F}} + v'_{2\mathrm{M}}. \tag{30}$$

The sign of $\mu'$ is not always fixed. But when $N$ is large enough (but finite), the absolute values of $v'_{1_\mathrm{F}}$ and $v'_{1M}$ are much larger than $v'_{2_\mathrm{F}}$ and $v'_{2\mathrm{M}}$. So the effect of $-v'_{2_\mathrm{F}} + v'_{2\mathrm{M}}$ is negligible, and $\mu' > 0$ almost always true. That means when mutant occurs in the sex with a higher dominance coefficient, natural selection is more efficient if the sex ratio of the population is biased towards this sex.

If we let $h_\mathrm{F} = 0$ and $h_\mathrm{M} = 1$, we have

$$\mu' = v'_{1\mathrm{M}} + v'_{2\mathrm{F}} - v'_{2\mathrm{M}}. \tag{31}$$

In this case, we find that $\mu' < 0$ almost always true. That is to say, when mutant occurs in the sex with a lower dominance coefficient, natural selection is more efficient if the sex ratio of the population is biased towards the other sex. The common conclusion drawn from the above two results is that when the dominance coefficients are different between two sexes, natural selection is more efficient in populations with sex ratios biased towards the sex with a larger dominance coefficient, no matter in which sex the mutant was originated.

### 4 Diffusion approximation

In this section, we use Kimura's diffusion approximation [5, 6] method to calculate the fixation probability of a single mutant in a diploid monoecious population.

#### 4.1 Population description and Hardy-Weinberg equilibrium

In a diploid monoecious population, there are $N$ individuals, each with two genetic sites. Each site is occupied by either A or B. Fitness and updating rules are consistent with previous sections.

There are three genotypes in the population, but Kimura's diffusion approximation method is only suitable for one-dimensional stochastic processes. Therefore, we adopt the Hardy-Weinberg equilibrium assumption of large-size populations [3]. Hardy-Weinberg equilibrium means that the fractions of alleles and genotypes satisfy:

$$\begin{cases} p_{\mathrm{AA}} = p_{\mathrm{A}}^2, \\ p_{\mathrm{BB}} = p_{\mathrm{B}}^2, \\ p_{\mathrm{AB}} = 2p_{\mathrm{A}}p_{\mathrm{B}}, \\ p_{\mathrm{A}} = p_{\mathrm{AA}} + \frac{1}{2}p_{\mathrm{AB}}, \\ p_{\mathrm{B}} = p_{\mathrm{BB}} + \frac{1}{2}p_{\mathrm{AB}}. \end{cases} \tag{32}$$

By doing so, we only need to focus on $p_{\mathrm{A}}$ and $p_{\mathrm{B}}$ instead of $p_{\mathrm{AA}}$, $p_{\mathrm{AB}}$, and $p_{\mathrm{BB}}$.

#### 4.2 Applying Kimura's diffusion approximation method to model allele frequency dynamics

The fixation probability of A when $p_{\mathrm{A}} = p$ can be expressed as

$$u[p] = \frac{\int_0^p G[x]dx}{\int_0^1 G[x]dx}, \tag{33}$$

where $G[x] = exp(-\int 2M_{\delta x}/V_{\delta x}dx)$. $M_{\delta x}$ and $V_{\delta x}$ are the mean change in allele frequency and the variance in that change over a generation, as functions of allele frequency $x$.

##### 4.2.1 Calculate $M_{\delta x}$ and $V_{\delta x}$

In our model, the number changes of $A$ can only be $+1$, $0$, or $-1$. So the changes of allele frequency $x$ and only be $+1/2N$, $0$, or $-1/2N$. We assume that $\alpha(x) = p(\Delta x = +\frac{1}{2N})$ and $\beta(x) = p(\Delta x = -\frac{1}{2N})$, and we have

$$\alpha(x) = x(1-x)\frac{x(1+\delta) + (1-x)(1+h\delta)}{x^2(1+\delta) + 2x(1-x)(1+h\delta) + (1-x)^2}, \tag{34}$$

$$\beta(x) = x(1-x)\frac{x(1+h\delta) + 1 - x}{x^2(1+\delta) + 2x(1-x)(1+h\delta) + (1-x)^2}. \tag{35}$$

Consequently, we have

$$M(x) = \frac{1}{2N}(\alpha(x) - \beta(x)), \tag{36}$$

$$V(x) = \left(\frac{1}{2N}\right)^2 (\alpha(x) + \beta(x)). \tag{37}$$

##### 4.2.2 When the mutant allele is recessive ($h = 0$)

When $h = 0$, we have

$$\alpha(x) = x(1-x)\frac{1 + x\delta}{x(1 + x\delta) + 1 - x}, \tag{38}$$

$$\beta(x) = x(1-x)\frac{1}{x(1 + x\delta) + 1 - x}. \tag{39}$$

Then we have

$$G[x] = e^{-4Nx}(\delta x + 2)^{\frac{8N}{\delta}}. \tag{40}$$

Then we have

$$\rho_A = u\left[\frac{1}{2N}\right] = \frac{\int_0^{\frac{1}{2N}} G[x]dx}{\int_0^1 G[x]dx} \approx \frac{1}{2N} + \frac{1}{6}\delta. \tag{41}$$

##### 4.2.3 When the mutant allele is dominant ($h = 1$)

When $h = 1$, we have

$$\alpha(x) = x(1-x)\frac{1 + \delta}{x(1 + \delta) + (1 - x)(1 - x\delta)}, \tag{42}$$

$$\beta(x) = x(1-x)\frac{1 + x\delta}{x(1 + \delta) + (1 - x)(1 - x\delta)}. \tag{43}$$

Then we have

$$G[x] = e^{4Nx}(x + \frac{\delta}{\delta + 2})^{-\frac{2N(4+4\delta)}{\delta}}. \tag{44}$$

Then we have

$$\rho_A = u\left[\frac{1}{2N}\right] = \frac{\int_0^{\frac{1}{2N}} G[x]dx}{\int_0^1 G[x]dx} \approx \frac{1}{2N} + \frac{1}{3}\delta. \tag{45}$$

#### 4.3 Results from the diffusion approximation are consistent with our approach

The results of diffusion approximation method are consistent with those of ancestor random walk method when $N \to \infty$ (the selection gradient is $1/6h + 1/6$).
